# Increased chromatin accessibility following 1α,25-dihydroxyvitamin D_3_ treatment in human endometrial stromal cells

**DOI:** 10.64898/2026.05.06.723064

**Authors:** MyeongJin Yi, Hamed Bostan, Francesco J. DeMayo

## Abstract

Vitamin D signaling has recognized roles in female reproductive physiology, but its effects at the chromatin level in endometrial stromal cells are still unclear. Here, we investigated how the active form of vitamin D, 1α,25-dihydroxyvitamin D_3_, or calcitriol, influences the accessible chromatin landscape of human endometrial stromal cells. Assay for transposase-accessible chromatin using sequencing (ATAC-seq) was performed on T-HESCs treated with either a vehicle or 1,25(OH)_2_D_3_. Ligand treatment increased overall chromatin accessibility, shown by higher ATAC-seq signal intensity, while causing only minor changes in the total number of called peaks. Peak annotation revealed that accessible regions were spread across both promoter-proximal and distal genomic areas. Integrating this data with CUT&RUN and RNA sequencing showed that most vitamin D-responsive cistromic modifications and transcripts were linked to nearby open chromatin, though fewer were associated with regions that were significantly differentially accessible. These results suggest that 1,25(OH)_2_D_3_-dependent transcription mainly occurs within a permissive, pre-accessible chromatin environment. This study offers new evidence that active vitamin D influences the epigenomic landscape of human endometrial stromal cells, establishing the chromatin-based molecular response to a chemically-defined VDR ligand, 1,25(OH)_2_D_3_, relevant to stromal differentiation and preparation for decidualization.

**Highlights:**

- First evidence suggesting the direct impact of active vitamin D, 1α,25-dihydroxyvitamin D_3_, 1,25(OH)_2_D_3_, enhanced the signal intensity of chromatin accessibility in human endometrial stromal cells
- Most accessible chromatin regions were shared between vehicle and ligand-treated human endometrial stromal cells
- 1,25(OH)_2_D_3_-responsive transcription occurs largely within pre-accessible chromatin in human endometrial stromal cells
- Assay for transposase-accessible chromatin sequencing (ATAC-seq) defines a chromatin-level pharmacologic response to a chemically defined VDR ligand in human endometrial stromal cells

## 1. Introduction

Vitamin D is known for its role in calcium homeostasis and skeletal health, but accumulating evidence indicates that it also contributes to reproductive physiology. In female reproductive tissues, vitamin D signaling has been implicated in endometrial receptivity, implantation, and decidual function (Yi *et al*., 2026), suggesting that this pathway influences the cellular environment required for pregnancy. Among its biologically active forms, 1α,25-dihydroxyvitamin D_3_, 1,25(OH)_2_D_3_, acts through the vitamin D receptor (VDR) (Haussler *et al*., 1968; Holick *et al*., 1971; Lawson *et al*., 1971), a ligand-activated nuclear receptor that regulates transcriptional programs involved in differentiation, inflammation, and hormone-responsive cellular processes (Haussler and Norman, 1969). As a chemically defined bioactive vitamin D metabolite and pharmacologic VDR ligand, 1,25(OH)_2_D_3_ provides an important model for understanding how nuclear receptor activation translates chemical exposure into molecular and cellular responses. Despite increasing interest in vitamin D action in the uterus, the chromatin-level effects of ligand exposure in human endometrial stromal cells remain insufficiently defined.

Endometrial stromal cells are highly dynamic and undergo functional reprogramming in response to steroid hormones and local signaling cues (Dunn *et al*., 2003). This plasticity requires coordinated regulation of gene expression, which is strongly shaped by chromatin state (Vrljicak *et al*., 2018; Vrljicak *et al*., 2023). Chromatin accessibility determines whether transcription factors and cofactors can engage regulatory elements such as promoters and enhancers (Thurman *et al*., 2012; Eustermann *et al*., 2024). Therefore, defining accessible chromatin landscapes provides mechanistic insight into how hormonal and metabolic signals are translated into transcriptional outputs. In the endometrium, where stromal cells undergo cyclical changes in proliferation, differentiation, and immune adaptation, chromatin accessibility is especially relevant for understanding how regulatory programs are established and modified (Gellersen and Brosens, 2014; Vrljicak *et al*., 2018; Vrljicak *et al*., 2023).

Assay for transposase-accessible chromatin using sequencing (ATAC-seq) is a powerful approach for interrogating genome-wide chromatin accessibility using relatively limited input material (Buenrostro *et al*., 2015; Miskimen *et al*., 2017). Compared with transcriptomic analysis alone, ATAC-seq provides information on the regulatory architecture that permits, constrains, or primes gene expression responses. This is particularly important in hormonally responsive cells, where transcriptional changes arise either through broad chromatin remodeling or through selective use of regulatory elements that are accessible (Wiench *et al*., 2011; Grontved and Hager, 2012; Hoffman *et al*., 2022; Gomez Acuna *et al*., 2024). From a pharmacologic perspective, defining chromatin accessibility provides a molecular mechanism for understanding how ligand exposure alters regulatory competence before or alongside measurable transcriptional output. Thus, ATAC-seq is well-suited to determine whether ligand treatment induces accessibility changes or instead acts within a pre-existing permissive chromatin landscape.

T-HESCs, an immortalized human endometrial stromal cell line, provide a useful model for investigating hormone-responsive signaling and chromatin regulation in the stromal compartment (Krikun *et al*., 2004a; Krikun *et al*., 2004b; Li *et al*., 2022). Because stromal cell differentiation is central to endometrial function and preparation for decidualization (Li *et al*., 2022), this system offers a biologically relevant context for examining how vitamin D influences the epigenomic architecture of uterine biology. Although previous studies have described transcriptional and functional effects of vitamin D in reproductive tissues, much less is known about whether 1,25(OH)_2_D_3_ alters chromatin accessibility in human endometrial stromal cells and how such changes relate to downstream gene expression.

In the present study, we used ATAC-seq to define the chromatin accessibility landscape of T-HESCs following treatment with 1,25(OH)_2_D_3_. We asked whether ligand exposure broadly remodels accessible chromatin or instead reinforces accessibility within an open regulatory environment. We further integrated chromatin accessibility profiles with transcriptomic data to assess the relationship between open chromatin regions and vitamin D-responsive genes. By focusing on the epigenomic consequences of exposure to a chemically defined VDR ligand, this study provides a concise pharmacologic scheme for understanding how active vitamin D influences regulatory architecture in human endometrial stromal cells.

## 2. Materials and methods

### 2.1. Materials and software

hTERT-immortalized endometrial stromal cells (T-HESCs) were purchased from the American Type Culture Collection (ATCC; Cat. CRL-4003, VA, USA). Dulbecco’s Modified Eagle Medium/Nutrient Mixture F-12 (DMEM/F12; Cat. 11330032), fetal bovine serum (FBS; Cat. 10082147), trypsin-EDTA (Cat. 25200056), optimized minimum essential medium (Opti-MEM; Cat. 31985070), and charcoal stripped FBS (Cat. 12676029) were purchased from Gibco (MA, USA). 1α,25-dihydroxy vitamin D_3_ (1,25(OH)_2_D_3_, calcitriol; Cat. 71820) was manufactured by Cayman (MI, USA). Penicillin-streptomycin was from Sigma-Aldrich (Cat. P0781, MO, USA). Qubit double-stranded (ds) DNA high sensitivity (HS) assay kit (Cat. Q32854), Qubit RNA high sensitivity (HS) assay kit (Cat. Q32852), TRIzol (Cat. 15596018), and UltraPure DEPC-treated water (Cat. 750023) were purchased from Invitrogen (CA, USA). Tagment DNA TDE1 Enzyme and Buffer Kits (Cat. 20034198; Illumina, USA) were used to prepare the ATAC-seq libraries. MinElute PCR purification kit was from Qiagen (Cat. 28004; Germany). Ingenuity Pathway Analysis was developed by Qiagen (CA, USA). RStudio-2025.05.0-496 and R 4.5.0 were from Posit PBC (MA, USA). The Integrative Genomics Viewer (IGV; University of California and Broad Institute, https://igv.org/app/) was used to load the peak track and visualize the signal.

### 2.2. Cell culture of human telomerase reverse transcriptase (hTERT)-immortalized human endometrial stromal cell line (T-HESC) and exposure of 1,25(OH)_2_D_3_ in vitro assays

The cells were cultured and maintained in Dulbecco’s Modified Eagle Medium/Nutrient Mixture F-12 (DMEM/F12), which was supplemented with 10% (v/v) fetal bovine serum (FBS), 1 mM sodium pyruvate, 100 units/mL penicillin, and 100 µg/mL streptomycin at 37°C, containing 5% CO_2_ in a humidified atmosphere (Michalski *et al*., 2018; Li *et al*., 2022; Maurya *et al*., 2022; Montague Redecke *et al*., 2025; Montague Redecke *et al*., 2026; Yi *et al*., 2026). To evaluate the impact of ligand, T-HESCs were treated with 2 nM 1,25(OH)_2_D_3_. The ligand was diluted in Opti-MEM supplemented with 2% (v/v) charcoal-stripped FBS, 1 mM sodium pyruvate, 100 units/mL penicillin, and 100 µg/mL streptomycin to minimize background hormonal effects from standard FBS. Vehicle-treated cells were prepared under the same culture conditions and used as controls for comparison with 1,25(OH)_2_D_3_-treated cells.

### 2.3. ATAC-seq library preparation

ATAC-seq libraries were prepared as previously described with minor modifications (Buenrostro *et al*., 2013; Buenrostro *et al*., 2015). Three technical replicates were generated for each treatment condition, resulting in a total of six samples. Cells were collected and washed with cold PBS, followed by lysis in Greenleaf buffer (10 mM Tris-HCl, 10 mM NaCl, 3 mM MgCl₂, and 0.1% IGEPAL CA-630). Nuclei were isolated by incubation on ice for 5 min and centrifugation at 500 × g for 5 min at 4 °C. Nuclei were counted using a hemocytometer, and 50,000 nuclei per sample were used for transposition. For the transposition reaction, nuclei were resuspended in 50 µL of transposase reaction mix containing 25 µL TD buffer, 2.5 µL Tn5 transposase (Illumina Nextera DNA Library Preparation Kit), and 22.5 µL nuclease-free water. Samples were incubated at 37 °C for 15 min. Following transposition, DNA was purified using the MinElute PCR Purification Kit (Qiagen) according to the manufacturer’s instructions. Purified DNA was amplified by PCR using 10 µL of eluted DNA, indexed adapters, and PCR master mix. Libraries were amplified for a total of 15 cycles. Amplified libraries were purified using 150 µL SPRI beads (Beckman Coulter). Beads were incubated at room temperature for 15 min, followed by magnetic separation for 5 min. The supernatant was removed, and the beads were washed twice with 80% ethanol. After air-drying, DNA was eluted in 50 µL elution buffer (10 mM Tris-HCl, pH 8.0). Final libraries were quantified and prepared for high-throughput sequencing according to standard protocols.

### 2.4. Evaluation of chromatin accessibility by 1,25(OH)_2_D_3_ in T-HESC using ATAC-seq

Sequencing was performed on a NovaSeq (Illumina). Raw reads (50 bp, paired-end) processed by adaptor trimming and quality filtering using Trim Galore version 6.7, with reads retained when the average quality score was greater than 20. Filtered reads were aligned to the human reference genome hg38 using Bowtie2 with unique mapping and up to two mismatches permitted for each read (-m 1 -v 2). Reads mapped to mitochondrial DNA and duplicated reads were removed before downstream analysis. Uniquely mapped, non-duplicated reads from each sample were normalized by down-sampling to 100 million reads. The first 9 bp of each read were used for downstream accessibility analysis.

Open chromatin regions were identified using MACS3 with a Q-value cutoff of 0.0001, followed by merging genomic intervals within 100 bp of each other (Zhang et al., 2008; Liu, 2014). Peaks identified from vehicle- and 1,25(OH)_2_D_3_-treated samples were used to evaluate global chromatin accessibility patterns, treatment-associated changes in ATAC-seq signal intensity, and differentially accessible regions (DARs). DARs were defined by comparing the normalized ATAC-seq signal between vehicle-and 1,25(OH)_2_D_3_-treated cells across merged peak intervals. (Zhang *et al*., 2008; Liu, 2014).

### 2.5. Annotation of the nearest gene and motif analysis from ATAC-seq called peaks

The gene associated with each ATAC-seq peak was predicted by searching for transcription start sites (TSSs) of nearby genes within a 100 kb range using the “annotatePeaks.pl function” of the Hypergeometric Optimization of Motif Enrichment (HOMER) motif discovery tool (Heinz *et al*., 2010; Roberson, 2018). Peak locations were annotated relative to genomic features, including promoter-proximal regions, untranslated regions, exons, introns, and intergenic regions. Promoter-associated peaks were defined based on their proximity to the TSS. HOMER’s findMotifsGenome.pl function was used to perform motif enrichment analysis for selected peak sets (Genomes Project *et al*., 2012; Khurana *et al*., 2013; Tuoresmaki *et al*., 2014; Wen *et al*., 2014)

For genomic feature classes, we assigned using transcript annotation and prioritized as promoter, 5′UTR, 3′UTR, exon, intron, or intergenic region, with promoter defined as within 2 kb of the nearest TSS. Reference gene annotation files were generated using the hg38 reference genome annotation. The hg38.refGene.gtf file was used for genomic feature annotation and downloaded from UCSC Genome Browser. “genePredToGtf -utr hg38 refGene hg38.RefGene.gtf” was used to generate transcript annotation files containing untranslated region information for downstream peak annotation.

### 2.6. Overlay of analyses of chromatin accessible peaks with cistromic and transcriptomic datasets

To compare accessible chromatin regions between vehicle- and 1,25(OH)_2_D_3_-treated T-HESCs, direct overlap analyses were performed using summit-centered ATAC-seq peak coordinates. Peak summit files for each condition were represented as 1 bp genomic positions, and each summit was extended by ±100 bp to generate fixed 200 bp windows centered on the peak summit. Overlap between vehicle and 1,25(OH)_2_D_3_ ATAC-seq peak sets was then assessed based on genomic intersection between these summit-centered windows. Peaks were classified as shared if a summit-centered window from one condition overlapped a summit-centered window from the other condition, and as condition-specific if no overlap was detected.

To assess concordance between accessible chromatin and ligand-associated cistromic regions, overlap analyses were also performed between 1,25(OH)_2_D_3_ ATAC-seq summit-centered windows and publicly available 1,25(OH)_2_D_3_ CUT&RUN peak intervals (GSE306127). ATAC-seq summits were extended by ±100 bp and intersected with CUT&RUN peak coordinates in hg38. Overlap was summarized as the number and proportion of ATAC summit windows that intersected CUT&RUN peaks, the number and proportion of CUT&RUN peaks that intersected ATAC summit windows, and the total number of overlapping peak pairs. Overlapping CUT&RUN peaks were further annotated to nearby transcripts and genes using the same nearest-TSS framework described above. Because these analyses were based on genomic proximity and coordinate intersection, overlap between datasets was interpreted as colocalization or association rather than direct evidence of regulatory interaction.

To relate accessible chromatin regions to transcriptomic responses, summit-centered ATAC-seq peaks or merged peak sets were annotated to nearby genes using the hg38 reference transcript annotation. The nearest transcription start site (TSS) was assigned to each peak, and peak-associated genes were compared with the list of differentially expressed genes (DEGs) identified from publicly available RNA-seq analysis of 1,25(OH)_2_D_3_-treated versus vehicle-treated T-HESCs (GSE254251). Peak-to-gene relationships were summarized both at the peak level and as collapsed lists of unique genes.

All coordinate parsing, overlap classification, nearest-transcript assignment, and summary table generation were performed computationally in Python 3.13. All transcriptomic regulators and pathways were analyzed bioinformatically in Qiagen IPA.

### 2.7. Validation of differentially accessible regions in open chromatin by qPCR

Total RNAs from T-HESCs were isolated using TRIzol reagent according to the manufacturer’s instructions. All isolated RNAs were quantified and certified using a Qubit RNA High Sensitivity kit (Invitrogen). Each cDNA was synthesized with a High-Capacity cDNA Reverse Transcription kit (Applied Biosystems), adding 2 µg of total RNA to the reaction mixture, and the reaction mixtures were incubated at room temperature for 10 min followed by additional incubation at 37°C for 2 hr. Each synthesized cDNA was quantified and certified using a Qubit dsDNA High Sensitivity kit (Invitrogen). The primers to amplify the enhancer region of GPAT3 (LOC112997542) (Barakat *et al*., 2018); forward 5’-GGGTCTTCAATAAACAGCAG-3’, reverse 5’-GTCACTGAGAACGACGTCTG-3’, and the enhancer regions of MAMDC2 (LOC127814909) (Barakat *et al*., 2018); forward 5’-GAATGGAATCAACTCGAGAG-3’, reverse 5’-GGAAGTCACGTAAATGAATG-3’, respectively. qPCR was performed with the CFX96 Real-Time PCR Detection System (Bio-Rad). Each value was derived from the comparative CT method, which compared the Ct value of one target gene to a reference gene using the 2^-ΔΔCt^ formula according to the manufacturer’s guidelines. ΔC_t_ indicates the differences in threshold cycles for target and reference (C_t, target_ – C_t, reference_), and ΔΔC_t_ represents the relative change in these differences between the target and reference (ΔC_t, target_ – ΔC_t, reference_). Therefore, the expression of the target, normalized to a housekeeping gene, was given by 2^-ΔΔCt^ and adjusted as a fold-change.

### 2.8. Statistical analysis

All quantitative data were first tested for normality. When the normality assumption was met, comparisons among multiple groups were performed using one-way ANOVA followed by Tukey’s multiple comparisons test, and comparisons between two groups were performed using Student’s t-test. For non-normally distributed data, comparisons among multiple groups were performed using the Kruskal-Wallis test followed by Dunn’s multiple comparisons test, and comparisons between two groups were performed using the Mann-Whitney test. Statistical significance was defined as *p* < 0.05 unless otherwise indicated. For sequencing-based analyses, normalized read counts and peak-associated signal intensities were used to evaluate differences between vehicle- and 1,25(OH)_2_D_3_-treated groups. Differential accessibility and transcriptomic overlap analyses were interpreted using adjusted statistical thresholds where applicable. Data were summarized as counts, percentages, or normalized signal intensity values, depending on the analysis.

## 3. Results

### 3.1. 1,25(OH)_2_D_3_ enhances chromatin accessibility in T-HESCs

To determine whether active vitamin D alters chromatin accessibility in human endometrial stromal cells, ATAC-seq was performed in T-HESCs treated with vehicle or 1,25(OH)_2_D_3_. Genome-wide ATAC-seq profiles revealed that ligand treatment enhanced chromatin accessibility, as reflected by increased ATAC-seq signal intensity across accessible regions in the maximum signal intensity at the midpoint of identified peaks (Fig. 1A) and open regions surrounding the transcriptional start sites (TSS) (Fig. 1B). Although ligand treatment enhanced ATAC-seq signal intensity, it did not substantially alter the overall peak landscape. Direct summit-centered overlap analysis revealed that most accessible regions were shared between vehicle- and 1,25(OH)_2_D_3_-treated cells, with 104,503 shared summits, corresponding to 80.85% of vehicle peaks and 90.15% of 1,25(OH)_2_D_3_ peaks (Fig. 1C and Supplementary Table 1). These findings indicate that 1,25(OH)_2_D_3_ primarily strengthens the accessibility of regulatory regions rather than inducing widespread chromatin opening. This result suggests that 1,25(OH)_2_D_3_ acts primarily within a pre-existing accessible chromatin landscape in T-HESCs, increasing the magnitude of accessibility at regulatory regions that are open rather than inducing widespread chromatin opening.

**Fig. 1.**
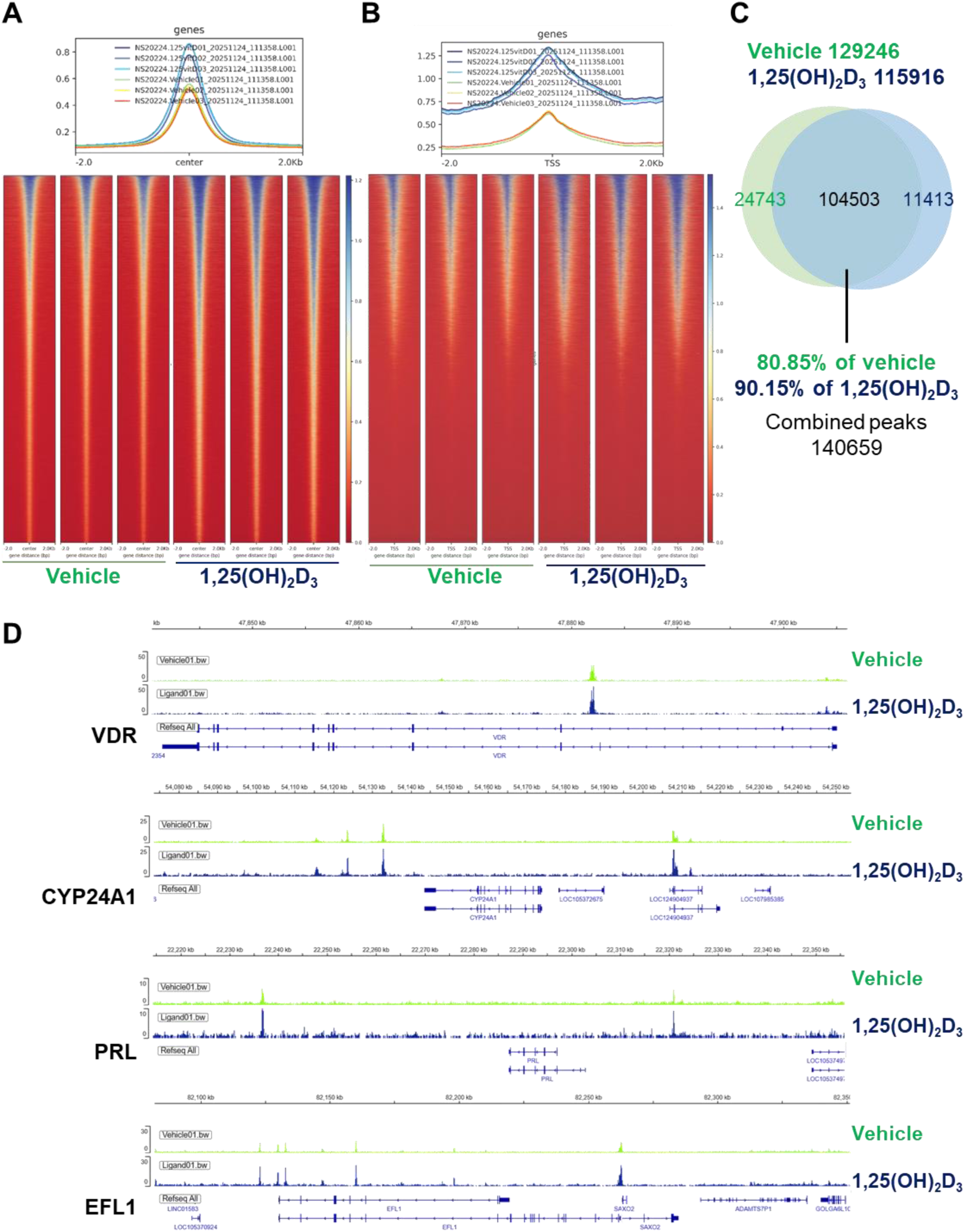
1,25(OH)_2_D_3_ enhances chromatin accessibility signal intensity in T-HESCs. The analysis shows that 1,25(OH)_2_D_3_ treatment is associated with increased accessibility signal without widespread reorganization of the global accessible chromatin landscape. (A) Overall profiles and heatmaps of ATAC-seq signal centered on called accessible regions (center ± 2 kb) and (B) surrounding transcription start sites (TSS ± 2 kb) in vehicle- and 1,25(OH)_2_D_3_-treated T-HESCs. (C) Overlap of open chromatin peaks identified by ATAC-seq in vehicle- and 1,25(OH)_2_D_3_-treated cells. (D) Representative genome browser loci of ATAC-seq signal at VDR, CYP24A1, PRL, and EFL1 regions from vehicle- and 1,25(OH)_2_D_3_-treated cells. RefSeq gene models are shown below.

### 3.2. 1,25(OH)_2_D_3_ preserves a largely shared accessible chromatin landscape

Although 1,25(OH)_2_D_3_ did not broadly increase the total number of accessible regions, comparison of called peaks between vehicle- and ligand-treated cells revealed condition-associated differences at the level of individual genomic intervals (Fig. 1D). Vehicle-treated cells contained 24,743 summit regions not overlapping the 1,25(OH)_2_D_3_ peak set, whereas 1,25(OH)_2_D_3_-treated cells contained 11,413 summits not overlapping the vehicle peak set (Fig. 1C and Supplementary Table 1). Thus, ligand treatment was associated with selective differences in peak representation, but these changes occurred within the context of a largely conserved accessible chromatin. These findings suggest that 1,25(OH)_2_D_3_ predominantly strengthens accessibility across a shared set of regulatory regions while introducing a more limited number of condition-associated accessible loci. This pattern supports a model in which ligand exposure modulates a permissive chromatin environment rather than establishing an entirely distinct accessibility architecture.

### 3.3. Distribution of differentially accessible regions (DARs) following ligand treatment

To further evaluate heterogeneity within ligand-responsive accessible regions, DARs were clustered according to ATAC-seq signal patterns across treatment conditions. Clustered heatmaps and average signal profiles revealed distinct groups of regions with treatment-associated accessibility changes (Fig. 2A). Notably, clusters showing higher signal in 1,25(OH)_2_D_3_-treated cells displayed stronger accessibility intensity compared with the corresponding vehicle condition, supporting the interpretation that ligand exposure enhances accessibility at selected chromatin regions. Together, these analyses indicate that 1,25(OH)_2_D_3_ produces a distinct set of DARs, with the most prominent effect reflected by increased signal intensity at a subset of ligand-responsive regions rather than by widespread gain of new accessible peaks. To identify genomic regions with statistically significant ligand-dependent changes in accessibility, differential accessibility analysis was performed between vehicle- and 1,25(OH)_2_D_3_-treated T-HESCs. Using an adjusted significance threshold and fold-change cutoff, we identified a set of DARs (Fig. 2B). Among these regions, 110 peaks showed increased accessibility following 1,25(OH)_2_D_3_ treatment, whereas 369 peaks showed reduced accessibility (Fig. 2B). Thus, the number of regions classified as decreased DARs was greater than the number of increased DARs. Annotation of DARs across genomic features showed that ligand-responsive accessibility changes were distributed across promoter-proximal and distal regulatory regions, including intronic and intergenic intervals (Fig. 2C). Increased DARs showed a relatively higher proportion of promoter-associated regions compared with decreased DARs. Intergenic regions represented the largest fraction of both DAR classes. This distribution suggests that 1,25(OH)_2_D_3_-responsive chromatin regulation involves both promoter-proximal and distal regulatory elements.

**Fig. 2.**
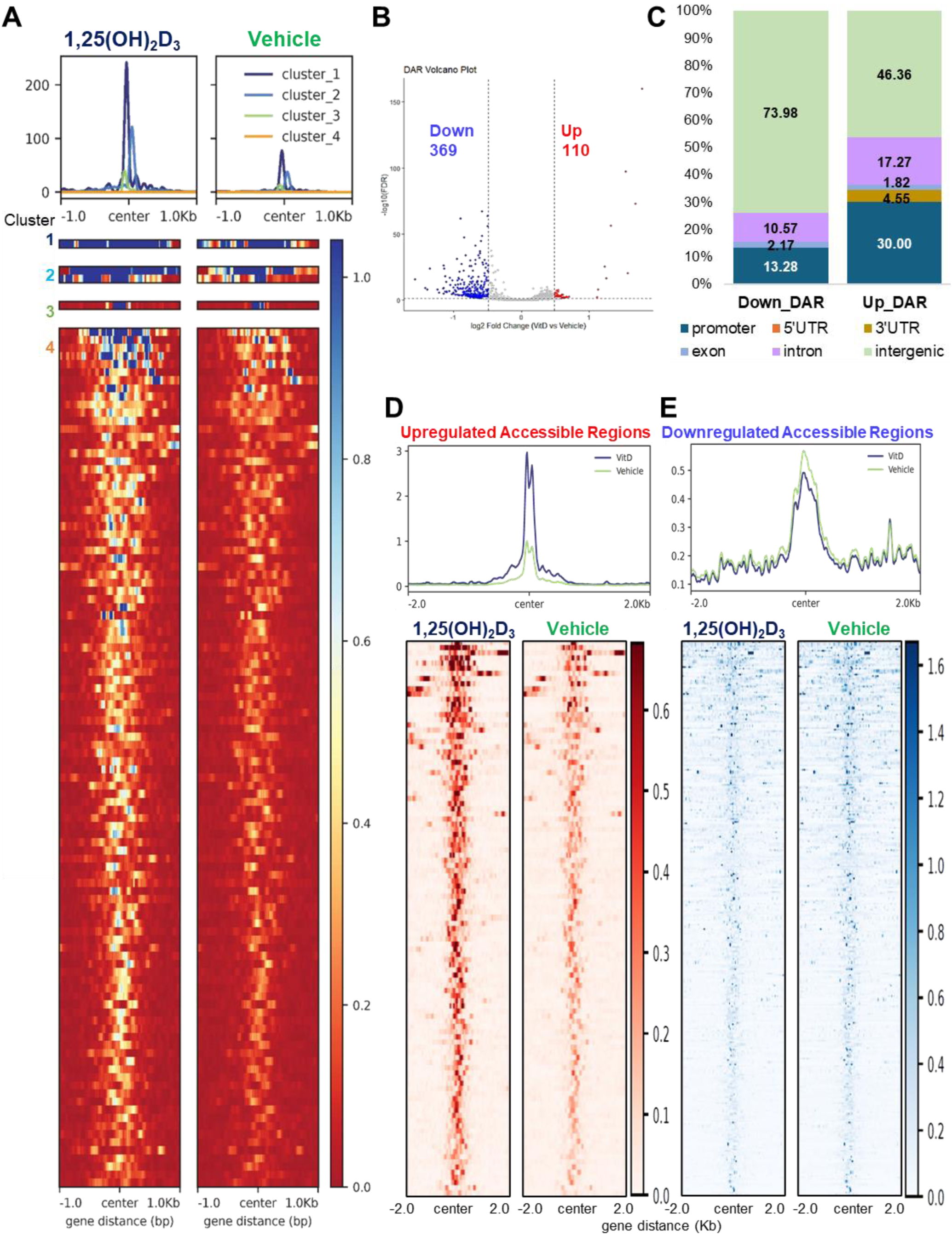
Differentially accessible regions (DARs) identified in response to 1,25(OH)_2_D_3_ in T-HESCs. (A) Clustered heatmaps and average signal profiles comparing ATAC-seq intensity patterns between vehicle- and 1,25(OH)_2_D_3_-treated cells across DARs. (B) Volcano plot showing differential chromatin accessibility between 1,25(OH)_2_D_3_-treated and vehicle-treated T-HESCs. Peaks were classified as increased or decreased accessible regions based on adjusted *p*-value and fold-change thresholds. (C) Comparison of genomic features of regions between loss and gain of accessibility. (D) Metaplot and heatmap of ATAC-seq signal across regions with increased accessibility in response to 1,25(OH)_2_D_3_, showing elevated signal intensity in ligand-treated cells compared with vehicle-treated cells. (E) Metaplot and heatmap of ATAC-seq signal across regions with decreased accessibility in response to 1,25(OH)_2_D_3_. The result shows that 1,25(OH)_2_D_3_ induces a distinct set of accessibility changes, with pronounced signal enhancement at selected ligand-responsive regions.

Because DAR counts alone do not fully capture the magnitude of accessibility changes, we next examined ATAC-seq signal intensity separately across increased and decreased DARs. Regions with increased accessibility showed a marked elevation of ATAC-seq signal in 1,25(OH)_2_D_3_-treated cells compared with vehicle-treated cells, as shown by both the average signal profile and heatmap visualization (Fig. 2D). This result indicates that, although increased DARs were fewer in number, they represented loci with clear ligand-enhanced accessibility. In contrast, regions classified as decreased DARs showed comparatively modest differences between treatment groups in both metaplot and heatmap analyses (Fig. 2E). These findings suggest that the accessibility gain induced by 1,25(OH)_2_D_3_ is concentrated at a subset of regulatory regions but is more pronounced in signal magnitude.

Overall, although increased DARs were fewer in number than decreased DARs, they showed a more pronounced gain in ATAC-seq signal intensity, indicating that 1,25(OH)_2_D_3_ enhances accessibility at a strongly responsive subset of regulatory regions. These results reinforce the model that active vitamin D modulates the T-HESC chromatin landscape primarily through quantitative strengthening of selected accessible regions, while the broader chromatin architecture remains largely pre-accessible.

### 3.4. Accessible chromatin regions overlap a subset of 1,25(OH)_2_D_3_-associated cistromic changes

To examine the relationship between accessible chromatin and ligand-associated cistromic changes, ATAC-seq summit-centered windows from 1,25(OH)_2_D_3_-treated T-HESCs were compared with publicly available 1,25(OH)_2_D_3_ CUT&RUN peak intervals (GSE306127). A total of 6,092 1,25(OH)_2_D_3_-associated CUT&RUN peaks from GSE306127 were included in the overlap analysis. Among these, 3,630 peaks overlapped accessible chromatin regions, representing 59.59% of the 1,25(OH)_2_D_3_-associated cistromic peaks (Fig. 3A and Supplementary Table 2). These findings indicate that a substantial proportion of ligand-associated cistromic changes occurs within regions of open chromatin.

**Fig. 3.**
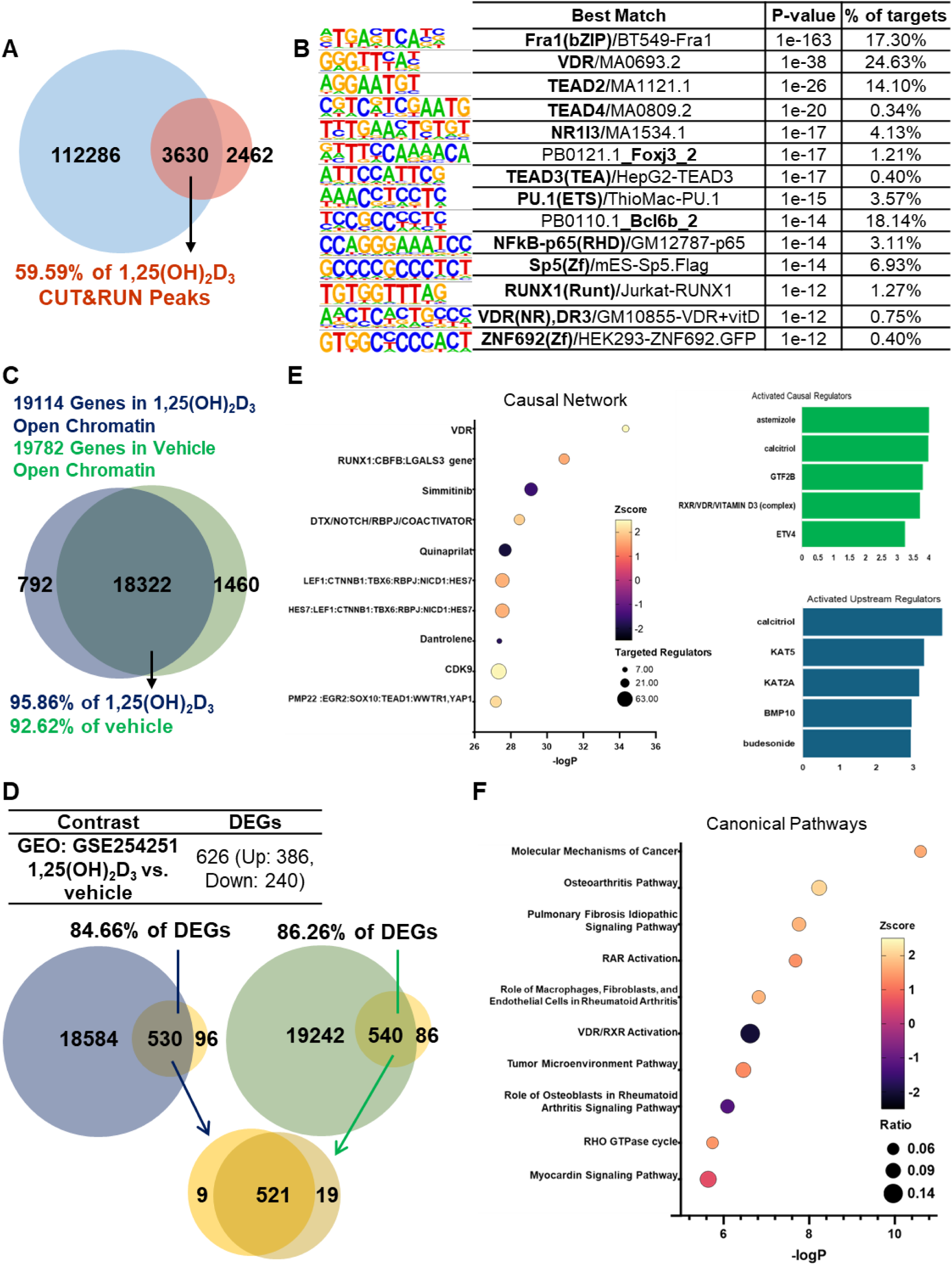
Integration of accessible chromatin-associated genes with vitamin D-responsive transcriptomic changes in T-HESCs. (A) The number of overlapped peaks between ATAC-seq and publicly available CUT&RUN. (B) Enriched binding motifs from cistromic modification peaks within the open chromatin areas in T-HESCs by HOMER *de novo* analysis. (C) Venn diagram showing overlap of genes annotated to ATAC-seq peaks in vehicle- and 1,25(OH)_2_D_3_-treated T-HESCs. Peaks were annotated to the nearest gene within 100 kb of the transcription start site. A total of 19,789 genes were associated with open chromatin regions in vehicle-treated cells, whereas 19,114 genes were associated with open chromatin regions in 1,25(OH)_2_D_3_-treated cells. Most annotated genes were shared between conditions, with 18,322 genes common to both datasets. (D) Overlap between 1,25(OH)_2_D_3_-responsive DEGs and genes associated with open chromatin regions in vehicle- or 1,25(OH)_2_D_3_-treated cells. Among 626 DEGs, 540 overlapped with vehicle-associated open chromatin genes, whereas 530 overlapped with 1,25(OH)_2_D_3_-associated open chromatin genes. The high proportion of overlap indicates that most ligand-responsive transcripts are associated with nearby accessible chromatin regions present in either condition. (E) Ingenuity Pathway Analysis (IPA) of the 530 differentially expressed genes associated with open chromatin regions in 1,25(OH)_2_D_3_-treated T-HESCs. Causal network analysis identified VDR as the most significant predicted activated master regulator. (F) IPA canonical pathway analysis of the same gene set revealed enrichment of pathways related to transcriptional regulation, immune-associated signaling, and vitamin D receptor activity.

To further characterize these shared regions, *de novo* motif analysis was performed using HOMER on the 3,630 overlapping peaks (Fig. 3B and Supplementary Table 2). The most significantly enriched motif corresponded to Fra1 (*p*-value = 1e-163; 17.30%), followed by motifs for VDR, TEAD2, TEAD4, NR1I3 (constitutive androstane receptor; CAR), FOXJ3, TEAD3, PU.1, NFκB, SP5, RUNX1, VDR (DR3), and ZNF692 (Fig. 3B and Supplementary Table 2). The enrichment of VDR-related motifs supports the ligand responsiveness of these shared regions, whereas the presence of additional transcription factor motifs suggests that 1,25(OH)_2_D_3_-associated cistromic regulation may occur in combination with other regulatory factors. Together, these results show that a subset of 1,25(OH)_2_D_3_-associated cistromic changes is localized within accessible chromatin regions and is enriched for motifs linked to ligand-responsive transcriptional regulation.

### 3.5. Accessible chromatin regions converge on a shared set of annotated genes

To assess whether ligand-associated differences in accessible chromatin altered the gene repertoire linked to open regulatory regions, ATAC-seq peaks were annotated to their nearest genes within 100 kb of the transcription start site (Supplementary Table 3). A total of 19,789 and 19,114 genes were associated with open chromatin regions in vehicle- and 1,25(OH)_2_D_3_-treated T-HESCs, respectively (Fig. 3C). The majority of annotated genes were shared between conditions, with 18,322 genes common to both datasets. These shared genes represented 92.62% of vehicle-associated genes and 95.86% of 1,25(OH)_2_D_3_-associated genes. These findings indicate that vehicle- and 1,25(OH)_2_D_3_-treated cells maintain a broadly similar open chromatin-associated gene repertoire, even though accessibility differs at selected genomic loci. Thus, ligand-dependent chromatin regulation appears to occur within an established stromal regulation rather than through wholesale reorganization of the genes associated with accessible chromatin.

### 3.6. Vitamin D-responsive transcripts are primarily associated with pre-existing accessible chromatin

To determine how chromatin accessibility relates to transcriptional responses to 1,25(OH)_2_D_3_, ATAC-seq peak-associated genes were integrated with RNA-seq data (GSE254251) generated from vehicle- and ligand-treated T-HESCs (Fig. 3D). Among the 626 differentially expressed genes (DEGs), 540 were associated with accessible chromatin regions in vehicle-treated cells, whereas 530 were associated with accessible chromatin regions in 1,25(OH)_2_D_3_-treated cells (Fig. 3D). A total of 521 DEGs were linked to accessible chromatin in both conditions, indicating that most ligand-responsive transcripts were associated with regulatory regions embedded within a shared accessible chromatin landscape (Supplementary Table 3). These findings suggest that transcriptional responses to 1,25(OH)_2_D_3_ occur predominantly within a permissive chromatin environment that is already accessible before ligand exposure, rather than through widespread establishment of newly accessible regions. Consistent with this interpretation, only a small subset of DEGs showed condition-specific associations with accessible chromatin regions. Specifically, 19 genes: COL10A1, CSF2RB, FABP4, FCMR, GLYAT, HCN2, KCNS2, LIF, LINC01081, LINC01123, MEST, MKX, PLEKHA7, PURG, RASSF2, STX1B, TGFB2, TNFRSF1B, and TRIM58 were uniquely associated with vehicle-specific accessible regions, whereas 9 genes: ARHGAP9, CCDC121, CYP3A5, CYTH4, NECAB1, NSDHL, PRRT4, SHANK1, and TUBE1 were uniquely associated with 1,25(OH)_2_D_3_-specific accessible regions (Supplementary Table 3). Thus, condition-specific chromatin associations accounted for only a minor fraction of ligand-responsive transcripts.

To further characterize the 1,25(OH)_2_D_3_-responsive transcripts associated with accessible chromatin, the 530 genes linked to open chromatin regions in 1,25(OH)_2_D_3_-treated cells were subjected to Ingenuity Pathway Analysis (IPA) (Fig. 3E and Supplementary Table 3). In causal network analysis, VDR was identified as the most significant predicted activated master regulator. Other highly ranked activating causal regulators included astemizole, calcitriol (1,25(OH)_2_D_3_), GTF2B, RXR/VDR/vitamin D_3_ complex, and ETV4 (Fig. 3E and Supplementary Table 3). In upstream regulator analysis, calcitriol, KAT5, KAT2A, BMP10, and budesonide were among the most strongly predicted activating regulators (Fig. 3E and Supplementary Table 3). Canonical pathway analysis showed enrichment of transcriptional and immune-related pathways, along with Molecular Mechanisms of Cancer, Osteoarthritis Pathway, Idiopathic Pulmonary Fibrosis Signaling Pathway, RAR Activation, Role of Macrophages, Fibroblasts and Endothelial Cells in Rheumatoid Arthritis, VDR/RXR Activation, Tumor Microenvironment Pathway, Role of Osteoblasts in Rheumatoid Arthritis Signaling Pathway, RHO GTPase Cycle, and Myocardin Signaling Pathway (Fig. 3E and Supplementary Table 3). These analyses indicate that transcripts associated with accessible chromatin in ligand-treated cells are enriched for regulatory networks centered on VDR and related transcriptional programs.

Overall, these findings indicate that 1,25(OH)_2_D_3_-responsive transcription in T-HESCs is largely associated with an established accessible chromatin framework. Rather than being accompanied by extensive generation of new open chromatin regions, ligand treatment appears to act mainly within pre-existing accessible regulatory regions linked to vitamin D-responsive gene expression.

### 3.7. Representative genomic loci support selective ligand-dependent chromatin remodeling

To further evaluate the relationship between ligand-responsive chromatin accessibility and transcriptional regulation, representative loci were selected from accessible regions overlapping 1,25(OH)_2_D_3_-responsive DEGs (Fig. 4A). Aggregate ATAC-seq profiles across ligand DEG-overlapping accessible regions further indicated that 1,25(OH)_2_D_3_-associated gain and loss peaks displayed distinct accessibility patterns between treatment conditions. The GPAT3-associated accessible region was derived from the upregulated accessible peak set overlapping ligand-responsive DEGs and showed increased accessibility after 1,25(OH)_2_D_3_ treatment in the ATAC-seq analysis (Fig. 4B). By contrast, the MAMDC2-associated accessible region was derived from the downregulated accessible peak set overlapping ligand-responsive DEGs and showed reduced accessibility following ligand treatment (Fig. 4B). Quantitative PCR analysis using RNA from vehicle- and 1,25(OH)_2_D_3_-treated T-HESCs confirmed ligand-associated regulation at these representative loci (Fig. 4C).

**Fig. 4.**
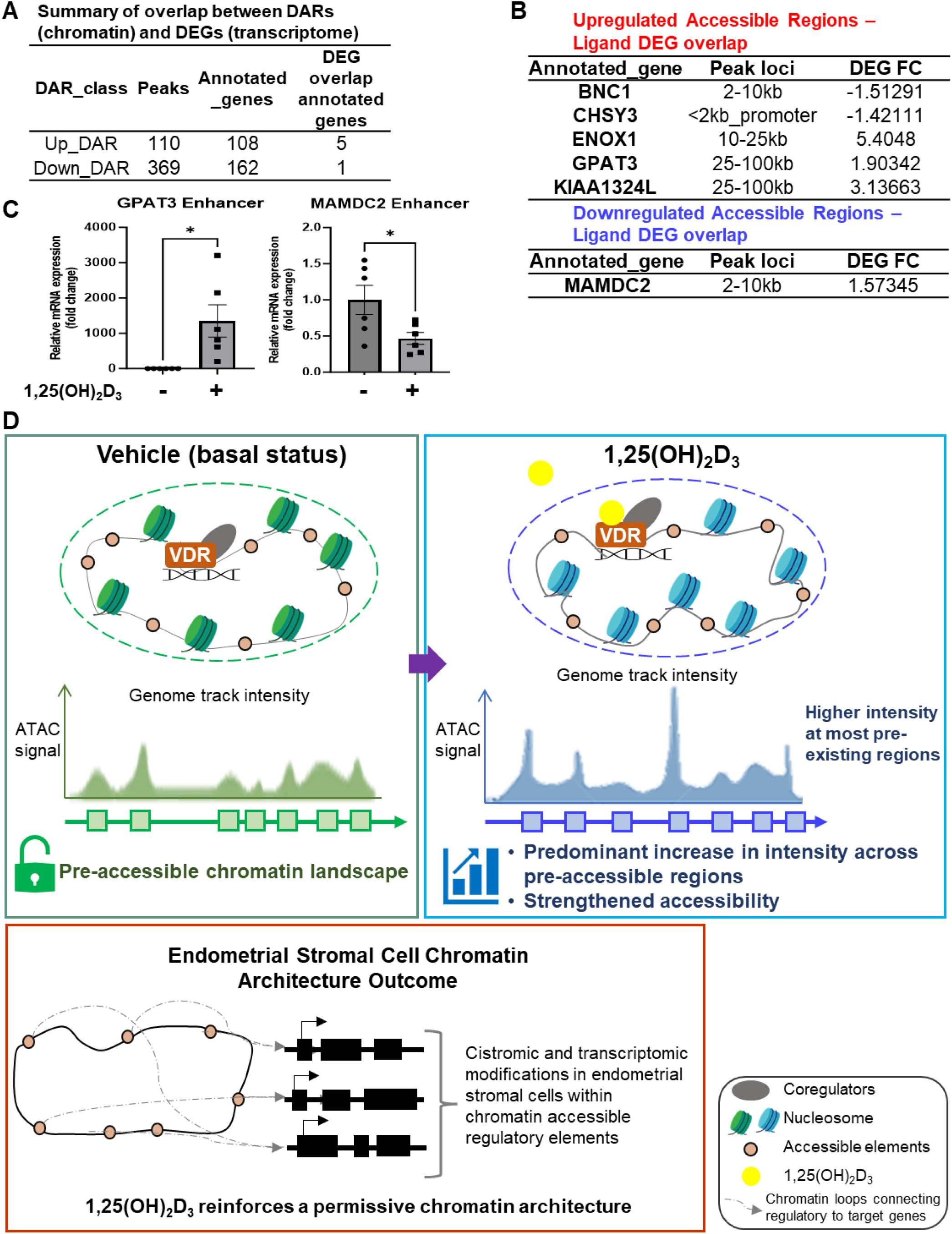
Schematic summary of selective chromatin remodeling and transcriptomic response to 1,25(OH)_2_D_3_ in T-HESCs. (A) Summarized transcript information on the overlap between DARs. (B) Upregulated DARs overlap with 1,25(OH)_2_D_3_ DEGs, and downregulated DARs overlap with 1,25(OH)_2_D_3_ DEGs. (C) Representative ligand-responsive accessible regions selected for locus-specific quantitative validation. Corresponding validation of the enhancer peak region of GPAT3 and MAMDC2. (D) Proposed working model of 1,25(OH)_2_D_3_-dependent chromatin-transcription coupling in T-HESCs. 1,25(OH)_2_D_3_ reinforces a permissive chromatin landscape in human endometrial stromal cells.

Collectively, these representative loci reinforce the genome-wide conclusion that active vitamin D acts primarily within a pre-existing accessible chromatin landscape in T-HESCs. Although 1,25(OH)_2_D_3_ induces selective changes at specific regulatory regions, the relationship between accessibility and transcription is locus-dependent and cannot always be inferred from the direction of change alone.

## 4. Discussion

This study provides new insight into how the active form of vitamin D, 1α,25-dihydroxyvitamin D₃, 1,25(OH)_2_D_3_, modulates chromatin accessibility in human endometrial stromal cells. From a pharmacologic perspective, 1,25(OH)_2_D_3_ is a chemically defined bioactive metabolite and ligand for VDR (Slatopolsky *et al*., 1984; Tsukamoto *et al*., 1991; Perez-Mijares *et al*., 1993), making it an appropriate model for investigating how nuclear receptor activation reshapes molecular regulatory architecture in target cells. Although vitamin D signaling has been studied in reproductive physiology, implantation-related biology, immune regulation, and decidual function (Hosseinirad *et al*., 2022; Rashidi *et al*., 2023; Metz *et al*., 2026), its direct impact on the accessible chromatin landscape of human endometrial stromal cells remains insufficiently defined. More broadly, chromatin accessibility itself has received relatively limited attention in endometrial stromal cell biology compared with transcriptional and hormonal regulation, despite its importance in determining which regulatory regions are available for signal-dependent gene control. By applying ATAC-seq to vehicle- and 1,25(OH)_2_D_3_-treated T-HESCs and integrating these data with transcriptomic profiles, we found that ligand exposure increased chromatin accessibility primarily at the level of ATAC-seq signal intensity, while most open chromatin regions remained shared between treatment conditions. Importantly, both a substantial subset of 1,25(OH)_2_D_3_-associated cistromic changes and most ligand-responsive transcripts were associated with regions within this accessible chromatin landscape. These findings indicate that active vitamin D does not broadly establish a new accessible chromatin landscape in this cellular context. Instead, 1,25(OH)_2_D_3_ appears to act largely within a pre-existing permissive chromatin environment, with selective accessibility changes at a smaller subset of loci.

A central observation of this study is that 1,25(OH)_2_D_3_ increased the magnitude of chromatin accessibility without causing a proportional expansion in the number of accessible regions. The most prominent effect of ligand treatment was enhanced ATAC-seq signal across regions that were accessible, rather than widespread formation of new peaks. This pattern supports a model in which active vitamin D strengthens the accessibility of pre-existing regulatory elements, potentially increasing their competence for transcription factor engagement or regulatory activity. This finding is particularly relevant to endometrial stromal cells, which must remain responsive to multiple hormonal, metabolic, inflammatory, and paracrine signals. Stromal cell differentiation and preparation for decidualization require extensive transcriptional coordination, yet these changes likely depend on a chromatin landscape that is poised or permissive before full differentiation occurs (Dunn *et al*., 2003; Krikun *et al*., 2004b; Gellersen and Brosens, 2014). In this context, vitamin D signaling functions more as a regulatory input that enhances the responsiveness of existing chromatin architecture. Such a mechanism would allow T-HESCs to adjust gene expression programs efficiently without requiring extensive chromatin reorganization (Li *et al*., 2022). This interpretation is consistent with the concept that stromal cells possess a flexible regulatory architecture that can be shaped by ligand-dependent nuclear receptor signaling (Michalski *et al*., 2018; Vrljicak *et al*., 2018).

The integration of ATAC-seq with RNA-seq further clarifies the nature of the ligand response in T-HESCs. Most 1,25(OH)_2_D_3_-responsive transcripts were associated with nearby accessible chromatin regions that were present in vehicle-treated cells, indicating that vitamin D-dependent transcription occurs largely within a pre-existing permissive regulatory environment. Similarly, overlap analysis with public CUT&RUN data showed that a substantial proportion of 1,25(OH)_2_D_3_-associated cistromic changes also occurred within open chromatin regions. Together, these observations indicate that both the cistromic and transcriptomic effects of 1,25(OH)_2_D_3_ are concentrated within chromatin regions that are already accessible. Few differentially expressed genes were linked specifically to differentially accessible regions, suggesting that broad transcriptional responsiveness to ligand exposure is not primarily driven by widespread creation of new accessible chromatin sites. Rather, the major contribution of 1,25(OH)_2_D_3_ may be to modulate the functional output of accessible regulatory elements. Thus, chromatin accessibility defines the regulatory territory available for ligand-responsive signaling, whereas ligand exposure influences how that territory is used. When peaks were annotated to nearby genes, the majority of associated genes were shared between conditions. This suggests that distinct accessible regions can converge on a broadly similar gene repertoire (Salamone *et al*., 2025). Gene regulation is rarely controlled by a single regulatory element; instead, promoters, enhancers, and distal regulatory regions often function as networks, with multiple elements contributing to the regulation of the same gene (Doane and Elemento, 2017; Vitale *et al*., 2021; Nazarova and Sexton, 2026). Therefore, a change in the accessibility of individual peaks does not necessarily imply a complete change in the associated gene program. The high degree of gene-level overlap observed in this study suggests that 1,25(OH)_2_D_3_ fine-tunes regulatory element usage within an established gene regulatory mechanism. This type of regulation could allow the same stromal gene repertoire to respond differently depending on ligand exposure, cellular state, or additional hormonal inputs.

The integration of ATAC-seq with RNA-seq further supports this interpretation. Most vitamin D-responsive transcripts were associated with nearby accessible chromatin regions (Fig. 3D). Thus, active vitamin D appears to act within an epigenomic landscape that is permissive for regulatory engagement. This suggests that the major contribution of 1,25(OH)_2_D_3_ is to increase the functional capacity or regulatory strength of accessible regions that are present. Ligand-activated transcription factors often regulate gene expression not only by binding to newly accessible DNA regions but also by redistributing among accessible regulatory elements, recruiting cofactors, altering local chromatin activity, or changing enhancer-promoter communication (Liu *et al*., 2014; Jin *et al*., 2025). In such cases, chromatin accessibility defines the regulatory territory available to the receptor, whereas ligand exposure determines how that territory is used. Although this study did not directly measure cofactor recruitment, histone modifications, or chromatin looping, the ATAC-seq patterns observed here are compatible with a model in which 1,25(OH)_2_D_3_ acts on an accessible chromatin landscape to modify transcriptional output. Future integration with VDR chromatin occupancy will be important to distinguish regions that VDR directly regulates from regions that change accessibility indirectly through downstream transcriptional or chromatin-associated mechanisms. These findings also have relevance for applied pharmacology because they illustrate how the action of a chemically defined ligand can be evaluated beyond conventional transcriptional endpoints (Pike, 2011; Nguyen *et al*., 2026). Pharmacologic responses are often measured through changes in gene expression, protein abundance, or cellular phenotype; however, ligand-induced changes in chromatin accessibility represent an earlier or parallel layer of molecular regulation (Chen *et al*., 2024). In this study, 1,25(OH)_2_D_3_ produced a prominent increase in ATAC-seq signal intensity without a corresponding large increase in peak number, suggesting that the pharmacological action of active vitamin D is expressed as a quantitative reinforcement of regulatory competence rather than broad chromatin remodeling. This distinction is important for understanding how bioactive metabolites and nuclear receptor ligands can modulate cellular state through changes in the magnitude of regulatory element accessibility (Etchegaray and Mostoslavsky, 2016).

Open chromatin provides a permissive state, but accessibility alone is not sufficient to determine whether a gene will be induced, repressed, or unchanged (Mansisidor and Risca, 2022). Many accessible regions remain functionally inactive under a given condition, while others become active only when appropriate transcription factors, cofactors, or signaling pathways are engaged (Chereji *et al*., 2019; Inge *et al*., 2024). Therefore, the presence of accessible chromatin should be viewed as a necessary but not always sufficient feature of transcriptional regulation. In T-HESCs, 1,25(OH)_2_D_3_ selectively activates or represses transcriptional programs by acting on a subset of accessible regions that are competent for vitamin D-responsive regulation. This interpretation is especially important for understanding stromal differentiation and preparation for decidualization. Decidualization requires the coordinated activation and repression of gene networks that support stromal cell transformation, extracellular matrix remodeling, immune communication, and endocrine responsiveness (Dunn *et al*., 2003; Krikun *et al*., 2004a; Gellersen and Brosens, 2014). Although the present study does not directly establish functional effects on pregnancy outcomes, implantation, or in vivo decidual success, it provides a chromatin-level framework for understanding how exposure to an active vitamin D alters the regulatory state of stromal cells. By increasing accessibility intensity across pre-existing regulatory regions, 1,25(OH)_2_D_3_ helps reinforce a chromatin architecture that supports differentiation-associated transcriptional responsiveness. This should be interpreted as a proposed chromatin-priming model rather than direct evidence of improved reproductive outcome (Fig. 4D).

If ligand treatment increases global ATAC-seq signal intensity, one might expect a large number of statistically significant DARs (Wang *et al*., 2022; Vu *et al*., 2023). Our results indicate analytical thresholds, peak-calling behavior, and the distinction between global signal shifts and region-specific statistical changes. Therefore, the modest number of DARs should not be interpreted as evidence that 1,25(OH)_2_D_3_ has little effect on chromatin. Rather, it indicates that the ligand’s effect is more strongly reflected in quantitative accessibility enhancement than in extensive gain or loss of discrete accessible regions (Fig. 4B and C). The genomic distribution of accessible regions and DARs further supports the idea that vitamin D signaling acts through both promoter-proximal and distal regulatory elements (Wang *et al*., 2023). Accessible chromatin regions were distributed across promoters, introns, exons, untranslated regions, and intergenic intervals, suggesting that ligand-responsive chromatin regulation is not confined to promoters alone (Wiench *et al*., 2011; Thurman *et al*., 2012). Distal intronic and intergenic regions include enhancers or other regulatory elements that contribute to stromal transcriptional programs (Heinz *et al*., 2010; Thurman *et al*., 2012). Thus, the gene associations reported here should be interpreted as active vitamin D acts primarily by reinforcing accessibility within a shared regulatory landscape, while producing selective remodeling at a smaller group of responsive loci (Fig. 4D).

Several limitations should be acknowledged. First, this study used T-HESCs as a model of human endometrial stromal cells. While T-HESCs are useful for mechanistic studies and retain important features of stromal biology, they do not fully recapitulate the complexity of primary stromal cells or the in vivo endometrial environment. Future studies using primary human endometrial stromal cells, decidualization models, or time-course designs would help determine whether the chromatin accessibility patterns observed here are maintained across more physiological contexts.

Second, this study was designed to define chromatin-level responses to a bioactive ligand rather than to evaluate reproductive toxicity, drug safety, or clinical pregnancy outcomes. Therefore, the findings should be interpreted as a mechanistic molecular pharmacology study of 1,25(OH)_2_D_3_ action in stromal cells. Future studies using primary human endometrial stromal cells, decidualization models, dose-response designs, or time-course experiments would help determine whether the chromatin accessibility patterns observed here are maintained across more physiology.

Third, ATAC-seq measures accessible chromatin but does not directly identify the transcription factors occupying these regions associated with regulatory activity, and integration with RNA-seq provides important correlative evidence but does not prove direct regulatory relationships between specific accessible regions and gene expression changes. Functional validation, such as perturbation of candidate regulatory elements, reporter assays, or CRISPR-based enhancer interrogation, would be required to establish causality.

Lastly, chromatin accessibility is only one layer of genome regulation. Three-dimensional genome organization, enhancer-promoter looping, nucleosome positioning, transcription factor cooperativity, and interactions with progesterone or estrogen signaling pathways are all likely to influence how vitamin D signaling is interpreted in stromal cells. These layers will be important to examine in future studies, particularly because endometrial stromal biology is shaped by multiple converging endocrine signals.

Despite these limitations, this study provides important insight into the epigenomic action of active vitamin D in human endometrial stromal cells. The accessible chromatin landscape of T-HESCs was largely shared between vehicle- and 1,25(OH)_2_D_3_-treated cells, indicating that ligand exposure acts primarily within an established permissive regulatory environment rather than by broadly creating a new one. 1,25(OH)_2_D_3_ enhanced ATAC-seq signal intensity across accessible regions and induced a subset of significant accessibility changes at selected loci. Integration with public cistromic data further showed that many 1,25(OH)_2_D_3_-associated chromatin occupancy changes occurred within open chromatin regions, while integration with RNA-seq data showed that most ligand-responsive transcripts were also associated with nearby accessible chromatin regions. Collectively, these observations support a model in which active vitamin D modulates transcription mainly through an accessible chromatin landscape, with both cistromic and transcriptomic responses occurring largely within open chromatin and with selective chromatin remodeling contributing at a smaller subset of loci. More broadly, these findings highlight chromatin accessibility as an important and relatively underexplored layer of regulation in human endometrial stromal cells.

In summary, this study identifies 1,25(OH)_2_D_3_ as a chemically defined modulator of chromatin accessibility in human endometrial stromal cells and highlights an understudied epigenomic dimension of vitamin D pharmacology in uterine biology. 1,25(OH)_2_D_3_ enhances accessibility intensity within an established regulatory landscape. This supports a model in which active vitamin D operates through a permissive stromal chromatin architecture that supports transcriptional responsiveness during stromal differentiation and preparation for decidualization. These findings also emphasize that chromatin accessibility is not merely a static backdrop, but an important regulatory framework in human endometrial stromal cells, a cell type in which this layer of regulation has received comparatively limited attention. These findings demonstrate the utility of chromatin accessibility profiling for defining molecular responses to bioactive ligands and provide a mechanism for future studies examining how vitamin D signaling interacts with VDR occupancy, three-dimensional genome organization, and ovarian steroid hormone pathways to regulate endometrial stromal cell function.

## CRediT authorship contribution statement

**MyeongJin Yi:** Conceptualization, Data curation, Formal analysis, Investigation, Methodology, Validation, Visualization, Writing – original draft, Writing – review & editing. **Bostan Hamed:** Data curation, Formal analysis, Software, Validation. **Francesco J. DeMayo:** Conceptualization, Funding acquisition, Resources, Supervision, Writing – original draft, Writing – review & editing.

## Acknowledgments

This research was supported in part by the Intramural Research Program of the National Institutes of Health (NIH). The contributions of the NIH author(s) were made as part of their official duties as NIH federal employees, are in compliance with agency policy requirements, and are considered Works of the United States Government. However, the conclusions presented in this paper are those of the author(s) and do not necessarily reflect the views of the NIH or the U.S. Department of Health and Human Services. The authors appreciate the support from Epigenomics and the DNA Sequencing Core (1ZICES102545), as well as the Integrative Bioinformatics Supportive Group (1ZICES103371).

## Appendix A. Supplementary material

Supplementary Table 1

Supplementary Table 2

Supplementary Table 3

## Supplementary data statement

Supplementary data associated with this article can be found in the online version.

## Data availability

The raw and processed data files generated in this study are available in NCBI GEO with the accession numbers: ATAC-seq (GSE326063). This study utilized publicly available RNA-seq data files (GSE254251) from the following samples: GSM8036787, GSM8036788, GSM8036789, GSM8036790, GSM8036791, and GSM8036792, and CUT&RUN data files (GSE306127).

The following datasets were generated:

| Contributor(s) | Year | Dataset title | Database and Identifier |
| --- | --- | --- | --- |
| Yi, MyeongJin; Bostan, Hamed; DeMayo, Francesco J | 2026 | Epigenomic changes in chromatin accessibility following 1,25-dihydroxyvitamin D<sub>3</sub> treatment in human endometrial stromal cells | GSE326063 |

